# Mitochondrial topoisomerase I (Top1MT) prevents the onset of metabolic dysfunction-associated steatohepatitis (MASH) in mice

**DOI:** 10.1101/2024.09.05.611454

**Authors:** SA Baechler, LK Saha, VM Factor, C Chitnis, A Dhall, D Becker, JU Marquardt, Y Pommier

## Abstract

High fat (HF) diet is a major factor in the development of metabolic dysfunction-associated steatotic liver disease (MASLD) and steatohepatis (MASH), and mitochondria have been proposed to play a role in the pathogenesis of HF diet-induced MASH. Because Mitochondrial topoisomerase I (Top1MT) is exclusively present in mitochondria and Top1MT knock-out mice are viable, we were able to assess the role of Top1MT in the development of MASH. We show that after 16 weeks of HF diet, mice lacking Top1MT are prone to the development of severe MASH characterized by liver steatosis, lobular inflammation and hepatocyte damage. Mice lacking Top1MT also show prominent mitochondrial dysfunction, ROS production and mitochondrial DNA (mtDNA) release, accompanied by hepatic inflammation and fibrosis. In summary, our study demonstrates the importance of Top1MT in sustaining hepatocyte functions and suppressing MASH.

## Introduction

Lifestyle changes have led to an alarming spread of obesity, type 2 diabetes and metabolic diseases. As a consequence, in the past decade, the increased prevalence of metabolic diseases has contributed to the burden of liver disease ^1^ with non-alcoholic fatty liver disease (NAFLD) affecting 30% of the adults globally and having high liver-related morbidity and mortality in the United States ^2^. NAFLD is currently referred to as metabolic dysfunction-associated steatotic liver disease (MASLD) ^3^, and patients with MASLD can progress to a severe form called metabolic dysfunction-associated steatohepatitis (MASH) ^2^, which was until recently referred to as nonalcoholic steatohepatitis (NASH) ^3^. A recent epidemiological study estimates the global prevalence of MASH to be approximately 5% ^2^.

MASH is characterized by steatosis, lobular inflammation, hepatocyte damage, and varying degree of fibrosis ^4^. It is a major risk factor for liver cirrhosis and hepatocellular carcinoma, an emerging cancer. Consequently, MASH is the second and fastest rising indication for liver transplant in the United States ^5^, which is largely associated with obesity ^1,6^. Given the epidemic spread of obesity, the number of patients with chronic liver disease is expected to increase dramatically in the next few years, putting an enormous economic burden on society ^7–9^. Therefore, understanding the molecular mechanisms leading MASH progression is essential for the development of effective preventive and treatment strategies.

The liver is the central metabolic organ responsible for supplying glucose, lipids and ketones to all peripheral tissues. Its abundant mitochondria are key metabolic hubs where oxidation of amino acids, pyruvate and fatty acids is coupled with respiratory chain activity and ATP production ^10^. Thus mitochondrial dysfunctions via dysregulation of mitochondrial metabolism have been proposed to play a key role in the pathogenesis of MASH ^11–13^. Although multiple mechanisms have been implicated in MASLD and MASH progression ^14^, the molecular mechanisms and key regulators underlying MASH pathogenesis and progession are still poorly understood, and mitochondrial dysfunctions have been observed in MASH ^10^.

The mitochondrial topoisomerase I (Top1MT) ^15^, exclusively present in mitochondria, is a critical factor for mtDNA metabolism and homeostasis ^16–20^. By releasing the mtDNA topological stress associated with transcription and replication, Top1MT is critical for mtDNA expansion and tissue regeneration ^18,21^. Our recent work also demonstrated that Top1MT also supports mitochondrial protein biosynthesis ^22^. As such, Top1MT protects mitochondrial function and regeneration in challenging microenvironments and allows cell proliferation in a nutrient-deprived and hypoxic environment ^18,21,22^. This led us to hypothesize that Top1MT might be essential for mitochondrial adaption during overnutrition with excessive dietary fat, by preventing hepatocyte damage and death.

## Results

### Lack of Top1MT promotes MASH in adult mice fed with high fat diet

To address whether Top1MT-deficient mice are more susceptible to MASH, we fed 8 weeks-old WT and Top1MT KO mice a high fat (HF) diet for 16 weeks (Supplementary Figure 1A). Food consumption and body weight were monitored three times a week. Although neither difference in food intake nor in body weight gain were observed between WT and Top1MT KO mice over time (Supplementary Figure 1B and C), some Top1MT-deficient mice showed reduced survival after consumption of high fat diet for 16 weeks (Figure 1A). In the surviving Top1MT KO mice, the liver-to body weight ratio was significantly increased at 16 weeks (Figure 1B), and liver damage was reflected as a greater elevation of alanine transaminase (ALT) levels in the serum of TopMT KO than WT mice (Figure 1C). Circulating serum cholesterol levels were differentially elevated in mice lacking Top1MT (Figure 1D), indicating metabolic impairment and diminished adaption to the diet change in Top1MT KO mice.

**Figure 1.**
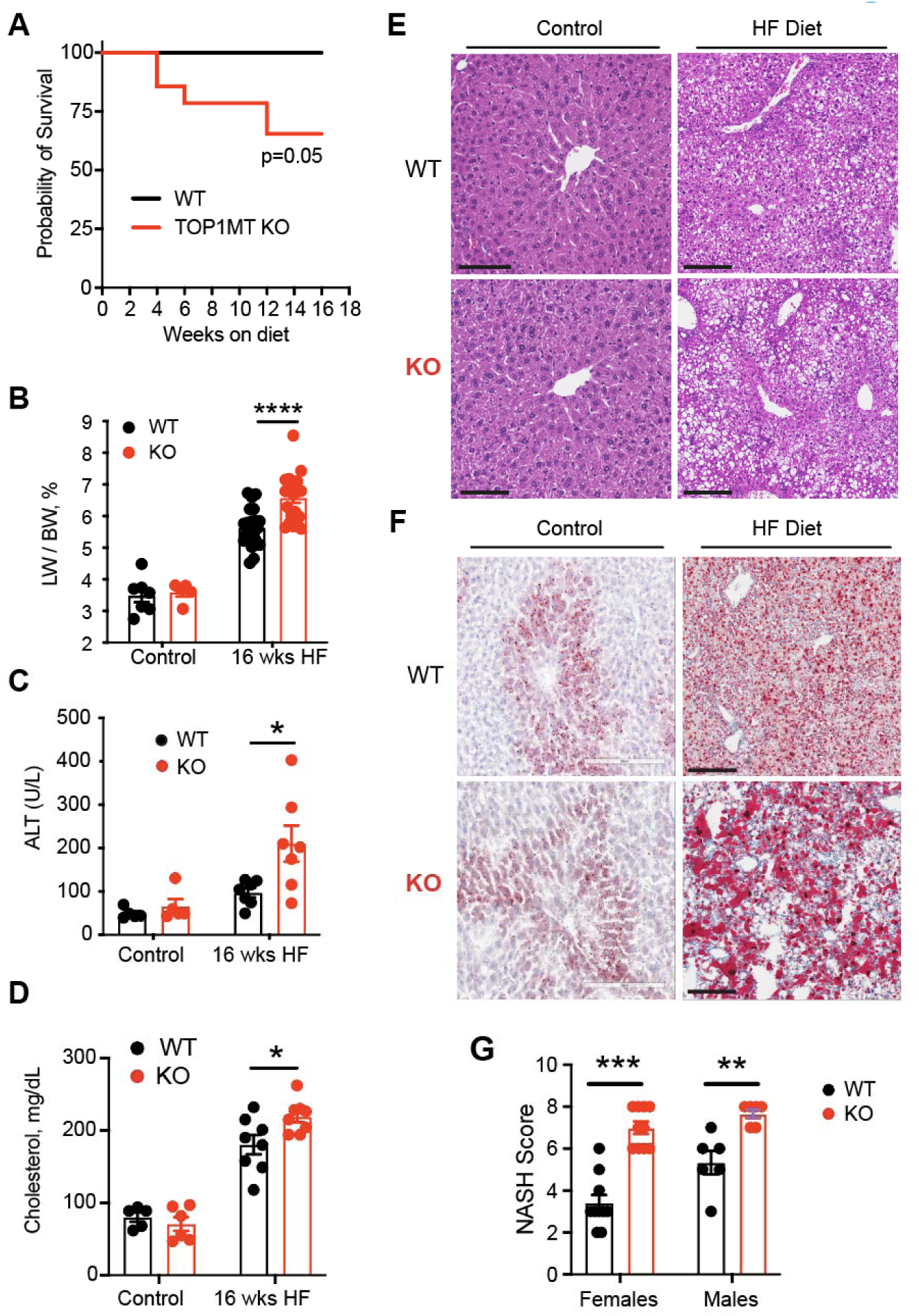
Top1MT deficiency promotes the onset of MASH in mice on a high fat diet (HF) for 16 weeks. **(A)** Kaplan-Meier survival curve of WT and Top1MT KO mice fed control or high fat (HF) diet (n=16). Gehan-Breslow-Wilcoxon test. **(B)** Liver to body weight ratio after control (WT: n=7; KO: n=5) or HF diet (WT:n=21; KO: n=21). **(C-D)** Plasma ALT and cholesterol levels after control. (WT: n=5; KO: n=5) or HF diet (WT: n=7; KO n=7). **(E)** Representative images of H&E (20x magnifications) staining of liver sections from mice fed control or HF diet.**(F)** Representative images of lipid staining using oil red O of liver sections. **(G)** MASH score quantified on liver histology (steatosis, ballooning, and inflammation). Data in bar graphs are presented as mean ± SEM. Statistical analysis was performed using a two-tailed Student’s t-test. Females (WT: n=10; KO: n=10) or Males (WT: n=5; KO n=5).

H & E staining of liver sections showed a marked increase in lobular inflammation and hepatocyte ballooning in the Top1MT KO animals fed with HF diet (Figure 1E). Histopathological analysis assessed by Oil Red O staining established pronounced steatosis in Top1MT KO mice with elevated lipid accumulation (Figure 1F). Assessment of the MASLD activity or MASH score further established enhanced disease progression in the Top1MT KO mice (Figure 1G). Notably, these features were more prominent in male animals, which is in line with previous reports that male C57Bl/6 are more susceptible to NASH development than their female counterparts ^23^. For this reason, we conducted the following cellular and molecular analysis in male animals only.

### The liver of *Top1MT* KO mice show prominent mitochondrial dysfunction in response to HF diet

Mitochondria play a crucial role in lipid metabolism and tend to associate with lipid droplets ^24,25^. Impaired mitochondrial functions have been shown in obese patients with MASH despite an increased mitochondrial mass ^26^. Using electron microscopy, we assessed mitochondria count and morphology in liver tissues. The number of mitochondria in contact with lipid droplets significantly increased in Top1MT deficient mice after 16 weeks on HF diet (Figure 2 A-B) while the overall mitochondrial count dropped slightly (Fig, 2C). The presence of two distinct mitochondrial populations possessing unique features and functions has recently been reported in brown adipose tissue, with peridroplet mitochondria (PDM) promoting lipid droplet expansion while fatty acid oxidation is largely carried out by cytoplasmic mitochondria (CM) ^27^. Beside an elevation in mitochondria-lipid droplets contact sites, the morphology of the mitochondria of Top1MT-deficient mice on a HF diet presented larger cross-sectional areas and an elongated shape compared to the WT controls (Figure 2D). This result is consistent with findings in brown adipose tissue showing that peridroplet mitochondria (PDM) are longer and larger in size ^27^. Mitochondrial DNA copy number was unchanged in liver samples of Top1MT-deficient mice compared to WT control (Figure 2E). The significant reduction in steady-state ATP levels in the livers of HF diet-fed Top1MT KO mice (Figure 2F) implies that the mitochondria of Top1MT KO mice are less functional under nutritional stress. Impairment of mitochondrial function is also consistent with elevated oxidative stress and lipid peroxidation levels in the liver of Top1MT-deficient mice after 16 weeks on a HF diet (Figure 2 G, I).

**Figure 2.**
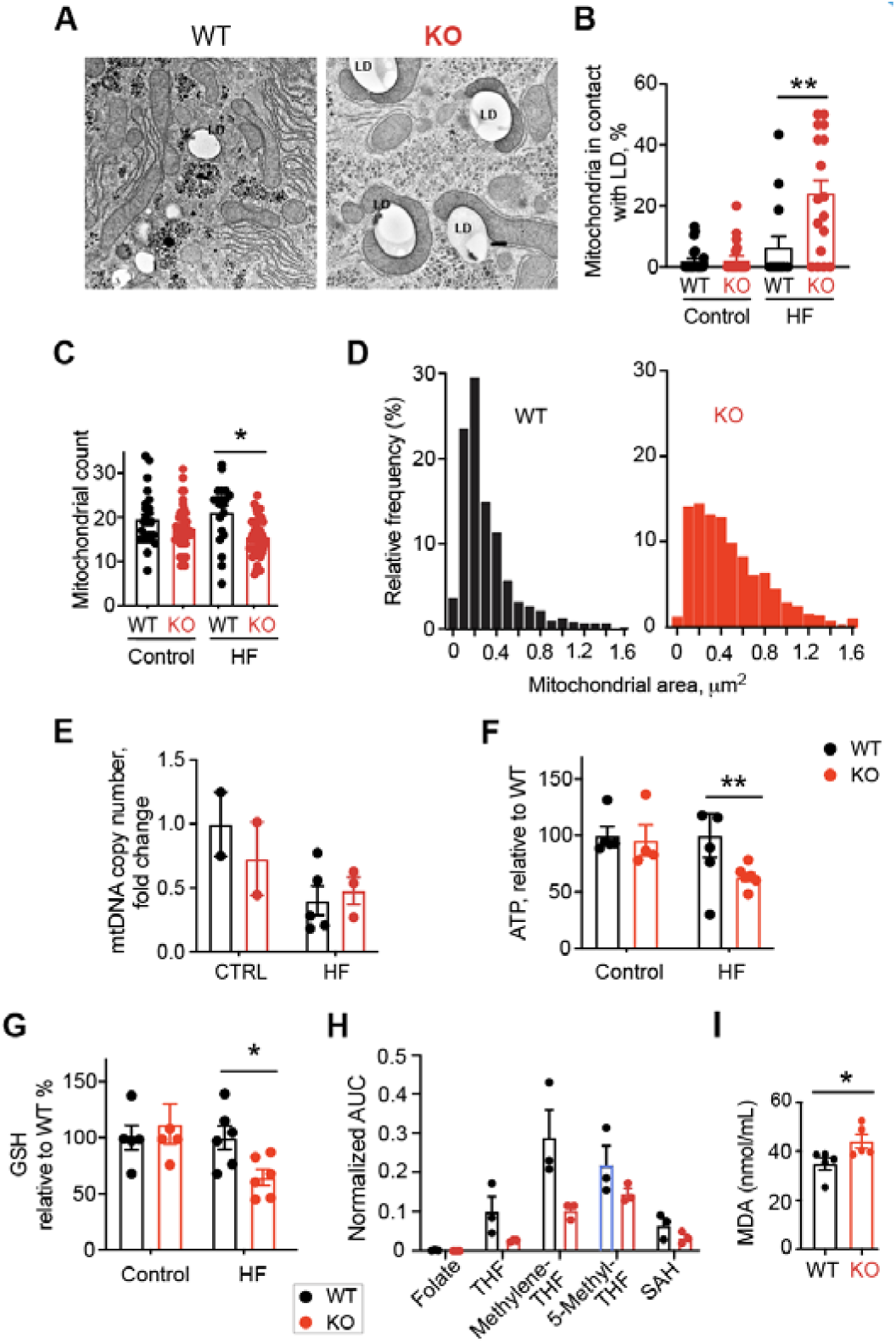
Top1MT deficiency causes mitochondrial dysfunction in mice on HF diet (16 weeks). **(A)** Representative electron micrographs of liver sections from WT and Top1MT KO mice after HF diet. Scale bar, 2 μm. **(B)** Number of mitochondria in contact with lipid droplets (LD) were quantified. n= number of electron micrograph per condition. Control (WT: n=24; KO: n=21). HF diet (WT:n=14; KO: n=17). **(C)** Number of total mitochondria after HF diet. n= number of electron micrograph per condition. Control (WT: n=24; KO: n=21) or HF diet (WT:n=20; KO: n=38). **(D)** Histogram showing increased mitochondrial perimeter in Top1MT KO mice after HF diet. **(E)** mtDNA copy number determined by RT PCR in liver tissue from WT and Top1MT KO mice. Control (WT: n=2; KO: n=2). HF diet (WT: n=5; KO n=3). **(F)** Cellular ATP content measured in liver tissue homogenate with ATPlite. Control (WT: n=5; KO: n=4). HF diet (WT: n=6; KO n=6). **(G)** Redox state determined by glutathione levels in the liver of WT and Top1MT KO. Control (WT: n=5; KO: n=4). HF diet (WT: n=6; KO n=6). **(H)** Metabolomic alterations in the folate cycle quantified by LC-MS in liver tissues. n=3 for both WT and Top1MT KO. **(I)** Redox state determined by malondialdehyde (MDA) levels in the liver of WT and Top1MT KO mice. n=3 for both WT and Top1MT KO.

Because alterations in folate cycle metabolism. i.e. folate deficiency, have been implicated in the development and progression of liver diseases including MASLD and MASH ^28^, we tested folate metabolites. Top1MT KO mice showed reduced folate metabolites, consistent with their defective mitochondrial functions in response to HF diet (Figure 2H). In addition, free fatty acids (FFA) triglycerides were significantly elevated in the plasma of Top1MT KO mice (Supplementary Figure 2A and B), indicative of alterations in glycolipid metabolism. Consistently, an increased expression of genes related to lipogenesis FASN and SREBP1 was observed in HF diet-fed mice lacking Top1MT (Supplementary Figure 2C). Together, these results demonstrate defective mitochondrial functions in the liver of Top1MT KO mice on HF diet.

### Mitochondrial abnormalities in the Top1MT hepatocytes isolated after one week of HF diet

To test the functionality of Top1MT-deficient hepatocytes, we performed *ex-vivo* cellular analyses (Figure 3A). HF diet significantly impacted the viability of hepatocytes, which had to be isolated after one week HF diet. Still, HF markedly reduced the viability yield to a greater extent in the Top11MT KO than in the WT hepatocytes (Figure 3B). Biochemical analyses revealed a significant increase in neutral lipids in the Top1MT hepatocytes (Figure 3C), while mitochondrial mass and mtDNA remained comparable in WT and KO hepatocytes (Figure 3D, E). Seahorse Flux Analysis of hepatocytes isolated from Top1MT-deficient mice subjected to HF diet showed significantly reduction of basal and maximal respiratory rate (Figure 3F), which was associated with a significant drop in ATP levels (Figure 3G). Because we recently reported defective mitochondrial translation in Top1MT KO ^22,29^, we measured mitochondrial protein synthesis by incorporation of ^35^S-methionine ^22^. Top1MT deficiency resulted in a pronounced decrease in mitochondrial translation in hepatocytes isolated from mice exposed to HF diet (Figure 3H). Impairment of mitochondrial functionality in turns led to an increase in mitochondrial superoxide (Figure 3K). These results demonstrate that lack of Top1MT profoundly affects the viability and functionality of hepatocytes isolated from mice subjected to HF diet.

**Figure 3.**
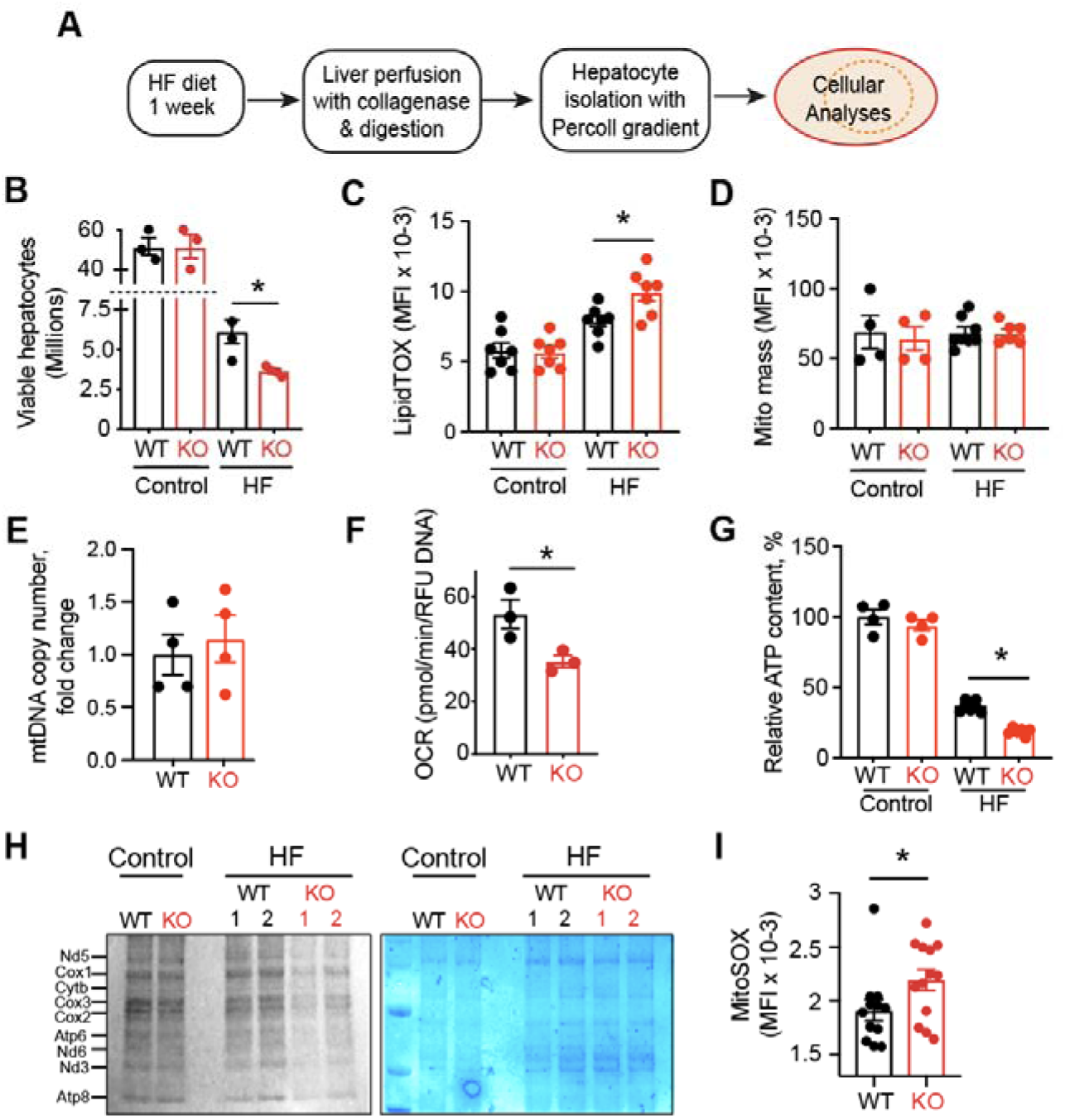
Mitochondrial abnormalities in primary hepatocytes isolated after HF diet for 1 week. **(A)** Experimental scheme. **(B)** Viable hepatocytes populations; n=3. Cell numbers were measured using trypan blue and a manual hemocytometer. Values are expressed as a percentage of viable cells. **(C)** Neutral lipid measurements: mean fluorescent intensity (MFI) of flow cytometry analysis after LipidTOX (H) (n=7) hepatocyte staining. Control (WT: n=7; KO: n=7) or HF diet (WT:n=7; KO: n=7). **(D)** Mitochondrial mass: mean fluorescent intensity (MFI) of flow cytometry analysis after MitoTracker Deep Red FM staining of hepatocytes. Control (WT: n=4; KO: n=4). HF diet (WT:n=8; KO: n=5). **(E)** mtDNA copy number determined by RT-PCR in hepatocytes. n=4 for both WT and Top1MT KO. **(F)** Oxygen consumption rates (OCR) measured by a Seahorse XF96 Extracellular Flux Analyzer. n=3 for both WT and Top1MT KO. Experiments were performed in quintuplets. **(G)** Relative ATP content of hepatocytes measured by ATPlite. Control (WT: n=4; KO: n=4) or HF diet (WT: n=7; KO n=6). (**H**) Mitochondrial protein synthesis measured by [^35^S]-methionine labeling of hepatocytes. A representative autoradiograph (left), and the corresponding loading control (comassie staining, right) is shown. **(I)** Mean fluorescent intensity of flow cytometry analysis after MitoSOX staining (CTRL: n=4; HF: n=6) of hepatocytes isolated after 1 week on a HF diet. n=12.

### Lack of Top1MT triggers hepatic inflammation and fibrosis and the release of mtDNA

The pathogenesis of MASLD/MASH is increasingly associated with enhanced innate immune responses ^30^. Plasma from MASH mice and patients contains high levels of circulating mtDNA, which can activate TLR9, and initiate an inflammatory response ^31^. Because of the observed hepatocyte ballooning together with the elevated ALT levels indicated increased hepatocyte damage in the liver of the Top1MT-deficient mice, we hypothesized that diminished mitochondrial translation might impair the assembly of oxidative phosphorylation complexes, leading to increased oxidative stress, lipid overload, and eventually to hepatocyte damage with the release of mtDNA into the blood. Accordingly, mtDNA plasma levels were significantly increased in the Top1MT-deficient mice under HF diet, while no difference was observed in circulating nuclear DNA (Figure 4A-C). Thus, the increase in circulating mtDNA parallels hepatic injury and elevated serum ALT levels ^31^.

**Figure 4.**
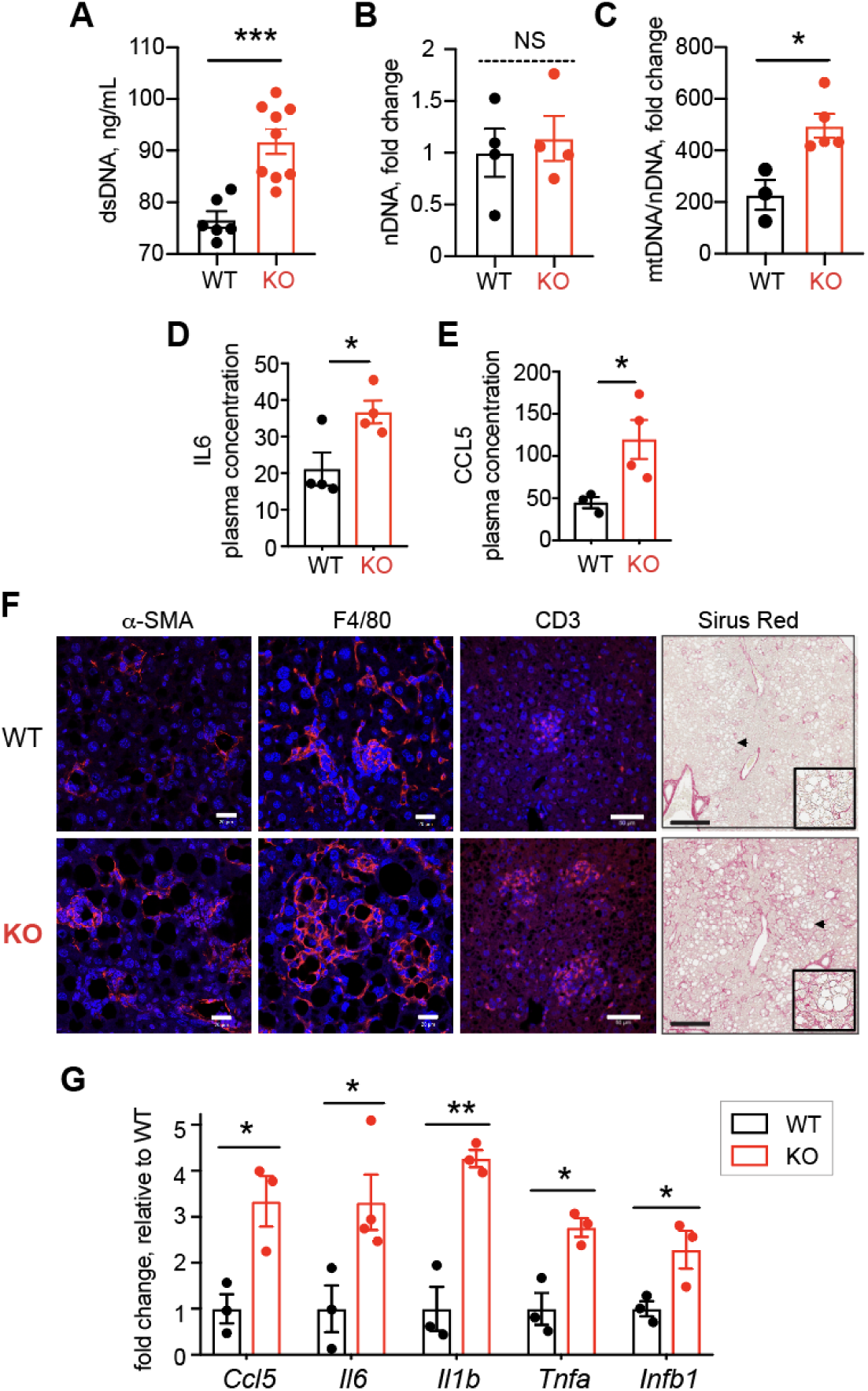
Top1MT deficiency triggers hepatic inflammation and fibrosis and mtDNA release in mice after 16 weeks HF diet. **(A)** Total dsDNA, nDNA, and mtDNA levels measured in the plasma of WT and Top1MT KO mice (WT: n=5; KO n=8). **(B)** Nuclear DNA (nDNA) were measured in the plasma of WT and Top1MT KO mice (WT: n=4; KO n=4). NS= not significant. **(C)** Mitochondrial DNA (mtDNA) levels in the plasma of WT and Top1MT KO mice. (WT: n=3; KO n=5). **(D)** Plasma levels of IL6 in WT and Top1MT KO mice (WT: n=4; KO n=4). **(E)** Plasma levels of CCL5 in WT and Top1MT KO mice (WT: n=3; KO n=4). **(F)** Representative images of Stellate cell (α-SMA), macrophage (F4/80) and T cell infiltration (CD3), were determined by immunofluorescence staining of liver tissue sections in WT and Top1MT KO mice fed a HF diet for 16 weeks. Scale bar, 20 μm. Representative images of Sirius red staining was used to measure collagen fiber deposition (right). Scale bar, 2 mm. **(G)** Transcript levels of proinflammatory genes; *Ccl5*, *Il6*, *IL1b*, *Tnfa*, and *Infb1* in liver tissues measured by RT-PCR.

Intra and extracellular mtDNA release activates inflammatory pathways with the activation of cytokines and innate immune cells ^12^. Correspondingly, plasma levels of the proinflammatory cytokines IL-6 and CCL5 were significantly elevated in the Top1MT-deficient mice on HF diet (Figure 4D-E). Presence of inflammation was further determined histologically by immunofluorescence staining for macrophages (F4/80), stellate cells (alpha-SMA) and T cells (anti-CD3) (Figure 4F). Enhanced infiltration of Kupffer cells, and alpha-SMA expression in stellate cells was found in liver sections of *Top1MT* KO mice (Figure 4F), consistent with recent findings of the activation of hepatic stellate cells in hepatic stress-induced progression of MASH ^30^. Additionally, a mild elevation of T-cell infiltrates was observed (Figure 4F). As extended periods of inflammatory response trigger the onset of fibrosis, we assessed the presence of liver collagen fibers by immunohistochemistry, staining connective tissue with Sirius Red. While no signs of fibrosis were present in WT liver sections of mice exposed to HF diet, Top1MT KO livers presented fibrosis (Figure 4F, right panel). Consistent with the inflammatory response, we found increased expression of a panel of proinflammatory cytokines in the Top1MT KO livers (Figure 4G). Together, these results demonstrate that functional Top1MT limits the release of mtDNA, and the inflammatory processes associated with MASH.

## Discussion

To date, dietary fat has been implicated in the progression of hepatic steatosis and MASH in both humans and animal studies ^32^. Although the exact cause of MASLD and MASH remains unclear, hepatic mitochondrial dysfunction have been proposed to be critical to the pathogenesis of MASH ^10^. In this study, we present the role of the mitochondria specific topoisomerase 1 (Top1MT) in reducing the development of MASH. By feeding a high fat diet (∼ 40% fat) to Top1MT KO mice, we show that Top1MT suppresses MASH development by preventing sustained mitochondrial dysfunction in hepatocytes and liver tissues (Figure 1-3). Our study also shows that lack of Top1MT aggravates a high fat diet-induced development of NASH associated with inflammation and fibrosis (Figure 4).

High fat diet with 40% calories coming from fat (mainly lard) was used in this study, which induced liver weight gain as expected (Fig. 1B), and Top1MT KO mice fed for 16 weeks with high fat diet developed hyperlipidemia and hypertransaminasemia (Figure 1C and D). In addition, high fat diet-fed Top1MT KO mice manifested a histological liver phenotype of MASH: lipid deposition (steatosis), local ballooning and liver damage (Figure. 1E-G). High fat diet in Top1MT KO mice eventually affects their lifespan (Figure 1A), recapitulating unhealthy lifestyle-associated MASH.

Many patients suffering from NASH have been reported to carry abnormal morphological changes in the hepatic mitochondria ^33^. Likewise, in our study, the assessment of mitochondrial content by transmission electron microscopy revealed increased mitochondrial area and decreased mitochondrial number in Top1MT KO mice than WT (Figure 2A-D). Compromised mitochondrial bioenergetic efficiency in hepatocytes of Top1MT KO is supported by their reduced OXPHOS capacity and ATP content compared to WT (Figure 2F, 3F-G). A reduction in the expression of mitochondrial protein assessed by [^35^S]-methionine incorporation revealed the lessening of mitochondrial quality control (Figure 3H). This is consistent with earlier reports that mitochondrial dysfunction and oxidative stress have been implicated in drug-induced MASLD progression in human liver cell lines ^34^. Increased ROS in liver tissue in Top1MT KO mice manifested as increased level of malondialdehyde (MDA) and decreased level of glutathione (GSH) (Figure 2G and I), in agreement with the possibility that NASH is a disease driven substantially by mitochondrial ROS overproduction. Increased presence of peridroplet mitochondria (PDM) in liver tissue (Figure 2A) and level of neutral lipids in hepatocytes (Figure 3C) of Top1MT KO mice suggests the dysregulated activation of lipogenesis in Top1MT KO mice. All these findings supports MASLD is a hepatic manifestation of the metabolic syndrome ^35^ and a ‘mitochondrial disorder’ ^36^. Mitochondrial stress including impaired mitochondrial DNA (mtDNA) homeostasis, dysfunction of mitochondria and altered metabolism lead to mtDNA release, which is increasingly associated with a broad range of diseases including liver inflammation ^37^. Release of mtDNA drives the generation of reactive oxygen species (ROS) and ROS-mediated mtDNA damage ^38^ with production of inflammatory cytokines including IL-6, IL1-β and TGF-α ^13^ and activation of the STING-cGAS pathway ^39^. Higher level of damaged mitochondria and ROS-mediated mtDNA damage have been found in human patients with MASH ^40,41^ and the release of damaged mtDNA into the plasma activates the endolysosomal toll-like receptor 9 (TLR9), causing pro-inflammatory responses ^31^. Indeed, we found increased mtDNA release, elevated gene expression and plasma level of proinflammatory molecules, and eventual hepatic inflammation in Top1MT KO mice. These findings corroborate the interdependency between mitochondrial dysfunction, ROS, mtDNA release and inflammation.

The ‘multiple hit ‘hypothesis including nutritional and genetic factors has been proposed for the development and progression of MASH ^42^. Indeed, previous reports have established the risk of MASLD and MASH with palatin-like phospholipase (PNPLA3) SNPs ^43^. Variants of transmembrane 6 superfamily member 2 (TM6SF2) gene also confer the susceptibility to MASLD ^44,45^, and it remains to be determined whether TOP1MT SNPs are present in MASH patients ^46^. To explore the potential clinical relevance of a *Top1MT* gene expression signature, we performed transcriptome analysis of WT and Top1MT-deficient liver samples after 16 weeks on a high fat diet. Supervised hierarchical clustering of 920 differentially expressed genes (Supplementary table 1) separated the murine liver samples into two clusters depending on Top1MT genotype (Supplementary Figure 3A). Pathway analysis showed enrichment of genes related PPARα activation and unfolded protein response, and further confirmed enhanced hepatic fibrosis and liver steatosis (Supplementary Figure 3C). Applying the *Top1MT* gene signature derived from the mice livers on the high fat diet to a cohort of MASH patients^47^ using integrative cluster analysis of orthologous genes separated the patients with mild NAFLD from the MASH patients (with advanced NAFLD) (Supplementary Figure 3B). Notably, Top1MT-deficient mice liver tissue clustered with the patients with MASH, while WT mice liver tissue clustered with patients with mild disease (Figure 5B). Thus, future investigations might use the Top1MT MASH signature as prognosis parameter. Additionally, systematic genomic studies might establish a link between human Top1MT variants ^39,46,48^ and the development and progression of NASH.

In conclusion, our study demonstrates that high fat diet-fed TOP1MT-deficient mice are highly susceptible to the development of MASH and have high hepatic inflammation in association with mitochondrial dysfunction, increased released of mtDNA and mtDNA-mediated formation of ROS. An understanding of the Top1MT-driven molecular pathways preventing MASH development may offer new strategies for therapeutic intervention.

## Acknowledgments

We are grateful to Michael Kruhlak (Head of Confocal Microscopy Core Facility, CCR, NCI, NIH) for support with confocal microscopy analyses and Dr. Jennifer Dwyer for scanning IHC slides. This study was supported by the Center for Cancer Research, the Intramural Program of the National Cancer Institute, National Institutes of Health, Bethesda, Maryland 20892 (Grant Z01 BC Z01 BC 006161-17 and Grant Z01 BC 006150-19 to Y.P).

## Conflict of interests

The authors declare no competing interests.

## Methods

### Animal experiments

As previously described ^19^ Top1MT knock-out (KO) mice were generated in C57BL/6 background and maintained by heterozygous breeding. All animal experiments followed according to the procedures outlined in the Guide for the Care and Use of Laboratory Animals under an animal study proposal approved by the NCI Animal Care and Use Committee. Mice were maintained at an American Association for the Accreditation of Laboratory Animal Care-accredited animal facility at the National Cancer Institute.

Mice were fed a normal chow diet (NIH31) for 8 weeks for proper growth and development. Thereafter, mice were randomly allocated to different experimental groups and fed a high fat (HF) diet (40.1% of the kcal derived from fat) and control diet (17% fat in the control diet), purchased from Dyets Inc. (Supplemental Figure 1A). Food was stored at −80^0^C and replaced three times a week to prevent fat oxidation, and food consumption and changes in body weight were recorded for the duration of treatment. The body and liver weight were recorded after the mice were euthanized. Mice were euthanized by cervical dislocation under 1% isoflurane anesthesia in accordance with approved protocol after one week and 16 weeks on diet for downstream analyses. Liver tissue and blood were collected for downstream analyses.

### Mice liver lipid and function measurements

Blood obtained by cardiac puncture under isoflurane anesthesia was centrifuged at 1000 g for 10 minutes at 4 °C. Plasma alanine aminotransferase (ALT), triglycerides (TG) and total cholesterol (TC) were measured with an automated biochemical analyzer (Sysmex CHEMIX-180, Japan) using commercial enzymatic assay kits (Applygen Technologies Inc., Beijing, China). All were evaluated in accordance with the manufacturer’s suggested methodology.

### Liver histology

Liver tissue was fixed overnight at 4°C in zinc formalin fixative (Sigma, #Z2902). Five micrometers paraffin sections were stained with hematoxylin and eosin (H&E) and Picro Sirius Red using standard techniques to assess liver morphology and degree of liver fibrosis. To assess hepatic steatosis, part of liver tissue was embedded in OCT, and frozen liver sections were stained with Oil Red O. The histological scoring of stained liver sections for steatosis grade, inflammation, hepatic fibrosis and hepatocyte ballooning was performed on H&E stained slides according to the NASH scoring system as previously described ^49^. Randomly selected fields were photographed using a Zeiss LSM 880 Microscope.

### Immunofluorescence staining of liver sections

Zink formalin-fixed, paraffin-embedded liver sections were immuno-stained with primary monoclonal antibodies against cell-specific markers of Stellate cells (mouse anti-α-SMA, 1:50, Dako), macrophages (rat anti-F4/80, 1:50, Bio-Rad Laboratories) and T cells (rabbit anti-CD3, Abcam Inc.) according to suppliers’ recommendation. Texas Red-conjugated anti-mouse and anti-rabbit were used as secondary antibodies. Images were taken and processed using Zeiss LSM 880 microscope.

### Transmission electron microscopy

Livers were fixed by in situ perfusion with 2% paraformaldehyde/2% glutaraldehyde in 1M cacodylate buffer (pH 7.4) at room temperature. Immediately after, tissue was cut into small fragments (1 x1 mm^3^) and post fixed in the same fixative for 2 hours at 4^0^C, followed by incubation in 1% osmium tetroxide for 1 h, then staining in 0.5% uranyl acetate for 1 h, and dehydration in a graded series of 35%, 50%, 70%, 100% (vol/vol) ethanol. Embedding in Epon and sectioning were performed as previously described ^21^. A combination of semi-thick (1 µm) and ultrathin (60 nm) sections was used to account for the metabolic zonation across hepatic sinusoids ^50^. Semi thick sections were stained with Toluidine Blue and used to outline the similar areas across hepatic lobule for the adequate comparison of mitochondria and lipid accumulation in WT and Top1MT KO livers. The ultrathin sections of the outlined areas were then prepared and imaged on an electron microscope operated at 80 kV (Hitachi H7650, Tokyo) by Electron Microscopy Core at the National Cancer Institute (Frederick, MD) as previously described ^21^. Digital images taken at x5000 magnification were captured by CCD camera (AMT, Danvers, MA) and used for analysis. Mitochondrial ultrastructure was analyzed with ImageJ software.

### Quantification of mtDNA copy number

Total DNA was isolated from liver tissues using PureLink Genomic DNA Mini Kit (Thermo Fisher Scientific, #K182001) following the manufacturers’ protocol. Quantitative PCR was performed in triplicates on 384-well reaction plates after DNA quantification with Nanodrop. For each PCR reaction, 25 ng DNA, 5 μL Power SYBR-Green PCR Master Mix (Applied Biosystems), and 0.5 μM of each forward and reverse primer were used. β2-microglobulin (β2m) was used as standard reference. Primer sequences used: β2m F (5′-TGCTGTCTCCATGTT TGATGTATCT-3′); β2m R (5′-TCTCTGCTCCCCACCTCTAAGT-3′); ND1 F (5′-AAGTCACCCTAGCCATCATTCTAC-3′); and ND1 R (5′-GCAGGAGTAA TCAGAGGTGTTCTT-3′).

### Glutathione (GSH) measurement

As previously described ^22^, the production of reactive oxygen species (ROS) was measured by quantifying glutathione (GSH) level in liver tissue. GSH levels were assessed in 50 mg liver tissue lysates using the luminescence-based GSH-Glo™ Glutathione Assay (Promega) according to the manufacturers’ protocol.

### Lipid peroxidation or Malondialdehyde (MDA) content detection

Lipid peroxidation was assessed using MDA assay kit (Cat# EEA015, Invitrogen) to measure the amount of MDA in liver tissue. The level of MDA was measured following the protocol provided by the manufacturer.

### LipidTOX assay

Assessment of lipid accumulation was done by staining mice hepatocytes with HCS LipidTOX™ Deep Red Neutral Lipid Stain (Cat# H34477) and detected by fluorescence microscopy according to the manufacturer’s protocol.

### Measurement of intracellular ATP levels

After homogenization of liver tissue (50 mg), ATP levels were determined using the ATPlite 1-step kit (PerkinElmer). Briefly, 100 μL ATPlite solution was added to 100 μL tissue homogenate in a 96-well white plate (Perkin Elmer Life Sciences, #6005680). After 5 min, luminescence was measured with an EnVision 2104 Multilabel Reader (PerkinElmer). Hepatocytes at a density of 5000 cells per well were seeded in 96-well plates and allowed to attach for 6 h. Cells were grown for 48 h in normal growth medium. Chemiluminescence was measured after adding 100 μL ATPlite solution (Perkin Elmer).

### Primary hepatocyte isolation

Hepatocytes were isolated by two-step collagenase perfusion of mouse liver followed by isodensity purification in Percoll gradient as previously described ^51^. The yield of viable cells was quantitated after staining with 0.4% trypan blue using automated cell counter (Cat# C10227, Invitrogen).

### Flow cytometry analysis of mitochondrial mass

After isolation of hepatocytes, cells (∼300,000) were seeded in 6-well plates and allowed to grow for 24 h. Cells were then incubated with 100 nM MitoTracker Deep Red dye (Thermo Fisher) for 20 min in normal culture medium at 37 °C. After three washes with 3 mL warm PBS, cells were trypsinized and resuspended in PBS supplemented with 1% FBS and 2 mM EDTA and immediately analyzed by flow cytometry (BD LSR Fortessa). Data analysis was performed using FlowJo v10 (FlowJo LLC).

### Measurement of oxygen consumption rate (OCR)

Isolated primary hepatocytes were seeded at a density of 20,000 cells/96-well. For OCR measurements, cells were incubated in medium supplemented with 2 mM glutamine, 10 mM glucose, and 1 mM pyruvate for 1 h, prior to the measurements using the XF Cell Mito Stress Kit (Agilent, #103015-100). For ECAR measurements (*n* = 3 for each genotype), cells were incubated with basal medium prior to injections using the Glycolysis Stress Test kit (Agilent, #103020-100). Experiments were run on a XF96 Extracellular Flux Analyzer (Seahorse Bioscience) in at least six replicates and raw data were normalized to DNA content measured using the CyQUANT Cell Proliferation Assay Kit (Thermo Fisher Scientific, #C7026).

### Mitochondrial translation

Mitochondrial protein synthesis in WT and Top1MT KO hepatocytes was assessed as previously described ^22^. In brief, WT and Top1MT KO hepatocytes were washed with methionine/cysteine-free DMEM, 2 mM L-glutamine, 96 μg/mL cysteine and 5% (v/v) dialyzed FBS and incubated for 10 min in the same media at 37 °C. 100 μg/mL emetine was then added directly to the media in order to inhibit cytosolic translation; after 20 min proteins newly synthetized in the mitochondrial compartment were labeled with 100 μCi [^35^S]-methionine (Perkin Elmer) for 1 h at 37 °C. Cells were then washed three times with PBS and lysed in PBS, 0.1% *n*-dodecyl-β-D-maltoside (DDM, Sigma), 1% SDS, 1X protease inhibitor cocktail (Roche) and 50 U benzonase (Novagen). 25 μg of total protein lysates were subjected to SDS-PAGE (Novex), gels were dried and radiolabelled proteins were detected by Phosphorimager.

### MitoSOX Red staining

The mitochondrial ROS levels were measured using a MitoSOX Red (Cat# HY-D1055, MedChemExpress, Shanghai, China) according to the manufacturer’s instructions. Isolated hepatocytes were seeded in 6-well plate, 24 h after plating cells were incubated with 5 μM MitoSOX Red solution for 30 min in dark at 37^0^ C with 5% CO2 incubator. Flow cytometry (BD LSR Fortessa) was performed immediately after trypsin digestion of cells.

### Free Fatty Acid (FFA) Quantitation

Hepatic FFA was measured in homogenized ∼25 mg liver tissue of WT and Top1MT KO mice by fluorescence detection using Free Fatty Acid Fluorometric Assay Kit (Cat# 700310, Cayman Chemical) according to the instructions from manufacturer.

### RNA extraction followed by quantitative PCR

Total RNA was extracted from liver tissue using TRIzol^TM^ (Thermo Fisher Scientific, #15596026) or RNeasy Mini Kit (Qiagen, #74104) and 1000 ng of total RNA reverse transcribed using RevertAid RT Reverse Transcription Kit (Thermo Fisher Scientific, #K1691) according to the manufacturers’ instructions. Real-time quantitative polymerase chain reaction (qPCR) was performed with the Applied Biosystems Vii7 using SYBR^®^ Green Master Mix. The primer sequences are shown in Supplementary Table 2. Gene expression was normalized to GAPDH or B2M (Thermo Fisher Scientific). The 2^-ΔΔCT^ method was used to analyze relative gene expression.

### RNA-Seq and clustering analysis

Total RNA was isolated from WT and Top1MT KO mice liver tissues of the same size (3 × 5 mm) using TRIzol^TM^ (Thermo Fisher Scientific, #15596026). Quality check (RIN ≥ 8) was performed with the Agilent Bioanalyzer. RNA-Seq libraries were prepared using the TruSeq Stranded RNA Prep Kit (Illumina, #RS-122-2201) and sequenced on the Ilumina HiSeq2500 instrument (125 bp paired end reads). Preprocessing and quality check of Illumina RNA-Seq data was performed using FastQC 0.10.1. followed by trimming of data by Trimmomatic-0.36. Reads were aligned against the reference genome (Mm 8) using BWA with default parameters. Only uniquely mapped reads were used for subsequent analyses followed by read summarization with featureCounts (subread-1.5.0-p1). All data analysis was performed using R programing language and related packages. The output matrix from featureCounts was input into the Bioconductor package DESeq2 for differential expression analysis. Significance testing was performed using R software (V.4.1.2) and Wald Test statistics. Genes were sorted based on their differences in peak intensity based on significant p-value and log2 fold changes of ±0.2. Functional networks were assessed using Ingenuity pathway analyses. The hierarchical clustering and visualization were created with the R package ComplexHeatmap (hclust) ^52^ using parameters “pearson” for distance and “complete” for cluster method.

### Statistical analyses

Statistical analyses were performed using GraphPad Prism v10 (GraphPad Software, Inc.). All data are represented as mean ± SEM or SD as indicated in the figure legends. The significant differences between groups were analyzed with Student’s t-test (parametric samples) and Wilcoxon rank-sum test (non-parametric samples). The degree of significance is indicated as *p < 0.05, **p < 0.01 and ***p < 0.001. NS= non-significant.

## Notes

### Competing Interest Statement

The authors have declared no competing interest.

## References

1. Hamid, O., Eltelbany, A., Mohammed, A., Alsabbagh Alchirazi, K., Trakroo, S., and Asaad, I. (2022). The epidemiology of non-alcoholic steatohepatitis (NASH) in the United States between 2010-2020: a population-based study. Ann Hepatol 27, 100727. 10.1016/j.aohep.2022.100727.

2. Younossi, Z.M., Golabi, P., Paik, J.M., Henry, A., Van Dongen, C., and Henry, L. (2023). The global epidemiology of nonalcoholic fatty liver disease (NAFLD) and nonalcoholic steatohepatitis (NASH): a systematic review. Hepatology 77, 1335–1347. 10.1097/HEP.0000000000000004.

3. Rinella, M.E., Lazarus, J.V., Ratziu, V., Francque, S.M., Sanyal, A.J., Kanwal, F., Romero, D., Abdelmalek, M.F., Anstee, Q.M., Arab, J.P., et al. (2023). A multisociety Delphi consensus statement on new fatty liver disease nomenclature. Hepatology 78, 1966–1986. 10.1097/HEP.0000000000000520.

4. Diehl, A.M., and Day, C. (2017). Cause, Pathogenesis, and Treatment of Nonalcoholic Steatohepatitis. N Engl J Med 377, 2063–2072. 10.1056/NEJMra1503519.

5. Wong, R.J., Aguilar, M., Cheung, R., Perumpail, R.B., Harrison, S.A., Younossi, Z.M., and Ahmed, A. (2015). Nonalcoholic steatohepatitis is the second leading etiology of liver disease among adults awaiting liver transplantation in the United States. Gastroenterology 148, 547–555. 10.1053/j.gastro.2014.11.039.

6. Cusi, K. (2012). Role of obesity and lipotoxicity in the development of nonalcoholic steatohepatitis: pathophysiology and clinical implications. Gastroenterology 142, 711–725 e716. 10.1053/j.gastro.2012.02.003.

7. Lazarus, J.V., Colombo, M., Cortez-Pinto, H., Huang, T.T.K., Miller, V., Ninburg, M., Schattenberg, J.M., Seim, L., Wong, V.W.S., and Zelber-Sagi, S. (2020). NAFLD — sounding the alarm on a silent epidemic. Nature Reviews Gastroenterology & Hepatology 17, 377–379. 10.1038/s41575-020-0315-7.

8. Berentzen, T.L., Gamborg, M., Holst, C., Sorensen, T.I., and Baker, J.L. (2014). Body mass index in childhood and adult risk of primary liver cancer. J Hepatol 60, 325–330. 10.1016/j.jhep.2013.09.015.

9. Younossi, Z.M., Tampi, R., Priyadarshini, M., Nader, F., Younossi, I.M., and Racila, A. (2019). Burden of Illness and Economic Model for Patients With Nonalcoholic Steatohepatitis in the United States. Hepatology 69, 564–572. 10.1002/hep.30254.

10. Fromenty, B., and Roden, M. (2023). Mitochondrial alterations in fatty liver diseases. J Hepatol 78, 415–429. 10.1016/j.jhep.2022.09.020.

11. Sunny, N.E., Bril, F., and Cusi, K. (2017). Mitochondrial Adaptation in Nonalcoholic Fatty Liver Disease: Novel Mechanisms and Treatment Strategies. Trends Endocrinol Metab 28, 250–260. 10.1016/j.tem.2016.11.006.

12. Prasun, P., Ginevic, I., and Oishi, K. (2021). Mitochondrial dysfunction in nonalcoholic fatty liver disease and alcohol related liver disease. Transl Gastroenterol Hepatol 6, 4. 10.21037/tgh-20-125.

13. Ramanathan, R., Ali, A.H., and Ibdah, J.A. (2022). Mitochondrial Dysfunction Plays Central Role in Nonalcoholic Fatty Liver Disease. Int J Mol Sci 23. 10.3390/ijms23137280.

14. Loomba, R., Friedman, S.L., and Shulman, G.I. (2021). Mechanisms and disease consequences of nonalcoholic fatty liver disease. Cell 184, 2537–2564. 10.1016/j.cell.2021.04.015.

15. Zhang, H., Barcelo, J.M., Lee, B., Kohlhagen, G., Zimonjic, D.B., Popescu, N.C., and Pommier, Y. (2001). Human mitochondrial topoisomerase I. Proc Natl Acad Sci U S A 98, 10608–10613. 10.1073/pnas.191321998.

16. Menger, K.E., Chapman, J., Diaz-Maldonado, H., Khazeem, M.M., Deen, D., Erdinc, D., Casement, J.W., Di Leo, V., Pyle, A., Rodriguez-Luis, A., et al. (2022). Two type I topoisomerases maintain DNA topology in human mitochondria. Nucleic Acids Res 50, 11154–11174. 10.1093/nar/gkac857.

17. Zhang, H., and Pommier, Y. (2008). Mitochondrial topoisomerase I sites in the regulatory D-loop region of mitochondrial DNA. Biochemistry 47, 11196–11203. 10.1021/bi800774b.

18. Khiati, S., Dalla Rosa, I., Sourbier, C., Ma, X., Rao, V.A., Neckers, L.M., Zhang, H., and Pommier, Y. (2014). Mitochondrial topoisomerase I (top1mt) is a novel limiting factor of Doxorubicin cardiotoxicity. Clin Cancer Res 20, 4873–4881. 10.1158/1078-0432.CCR-13-3373.

19. Zhang, H., Zhang, Y.W., Yasukawa, T., Dalla Rosa, I., Khiati, S., and Pommier, Y. (2014). Increased negative supercoiling of mtDNA in TOP1mt knockout mice and presence of topoisomerases IIalpha and IIbeta in vertebrate mitochondria. Nucleic Acids Res 42, 7259–7267. 10.1093/nar/gku384.

20. Douarre, C., Sourbier, C., Dalla Rosa, I., Brata Das, B., Redon, C.E., Zhang, H., Neckers, L., and Pommier, Y. (2012). Mitochondrial topoisomerase I is critical for mitochondrial integrity and cellular energy metabolism. PLoS One 7, e41094. 10.1371/journal.pone.0041094.

21. Khiati, S., Baechler, S.A., Factor, V.M., Zhang, H., Huang, S.Y., Dalla Rosa, I., Sourbier, C., Neckers, L., Thorgeirsson, S.S., and Pommier, Y. (2015). Lack of mitochondrial topoisomerase I (TOP1mt) impairs liver regeneration. Proc Natl Acad Sci U S A 112, 11282–11287. 10.1073/pnas.1511016112.

22. Baechler, S.A., Factor, V.M., Dalla Rosa, I., Ravji, A., Becker, D., Khiati, S., Miller Jenkins, L.M., Lang, M., Sourbier, C., Michaels, S.A., et al. (2019). The mitochondrial type IB topoisomerase drives mitochondrial translation and carcinogenesis. Nature Communications 10, 83. 10.1038/s41467-018-07922-3.

23. Matsushita, N., Hassanein, M.T., Martinez-Clemente, M., Lazaro, R., French, S.W., Xie, W., Lai, K., Karin, M., and Tsukamoto, H. (2017). Gender difference in NASH susceptibility: Roles of hepatocyte Ikkbeta and Sult1e1. PLoS One 12, e0181052. 10.1371/journal.pone.0181052.

24. Benador, I.Y., Veliova, M., Liesa, M., and Shirihai, O.S. (2019). Mitochondria Bound to Lipid Droplets: Where Mitochondrial Dynamics Regulate Lipid Storage and Utilization. Cell Metab 29, 827–835. 10.1016/j.cmet.2019.02.011.

25. Veliova, M., Petcherski, A., Liesa, M., and Shirihai, O.S. (2020). The biology of lipid droplet-bound mitochondria. Semin Cell Dev Biol 108, 55–64. 10.1016/j.semcdb.2020.04.013.

26. Koliaki, C., Szendroedi, J., Kaul, K., Jelenik, T., Nowotny, P., Jankowiak, F., Herder, C., Carstensen, M., Krausch, M., Knoefel, W.T., Schlensak, M., and Roden, M. (2015). Adaptation of hepatic mitochondrial function in humans with non-alcoholic fatty liver is lost in steatohepatitis. Cell Metab 21, 739–746. 10.1016/j.cmet.2015.04.004.

27. Benador, I.Y., Veliova, M., Mahdaviani, K., Petcherski, A., Wikstrom, J.D., Assali, E.A., Acin-Perez, R., Shum, M., Oliveira, M.F., Cinti, S., et al. (2018). Mitochondria Bound to Lipid Droplets Have Unique Bioenergetics, Composition, and Dynamics that Support Lipid Droplet Expansion. Cell Metab 27, 869–885.e866. 10.1016/j.cmet.2018.03.003.

28. Yang, M., Wang, D., Wang, X., Mei, J., and Gong, Q. (2024). Role of Folate in Liver Diseases. Nutrients 16. 10.3390/nu16121872.

29. Baechler, S.A., Dalla Rosa, I., Spinazzola, A., and Pommier, Y. (2019). Beyond the unwinding: role of TOP1MT in mitochondrial translation. Cell Cycle 18, 2377–2384. 10.1080/15384101.2019.1646563.

30. An, P., Wei, L.L., Zhao, S., Sverdlov, D.Y., Vaid, K.A., Miyamoto, M., Kuramitsu, K., Lai, M., and Popov, Y.V. (2020). Hepatocyte mitochondria-derived danger signals directly activate hepatic stellate cells and drive progression of liver fibrosis. Nat Commun 11, 2362. 10.1038/s41467-020-16092-0.

31. Garcia-Martinez, I., Santoro, N., Chen, Y., Hoque, R., Ouyang, X., Caprio, S., Shlomchik, M.J., Coffman, R.L., Candia, A., and Mehal, W.Z. (2016). Hepatocyte mitochondrial DNA drives nonalcoholic steatohepatitis by activation of TLR9. The Journal of Clinical Investigation 126, 859–864. 10.1172/JCI83885.

32. Lian, C.Y., Zhai, Z.Z., Li, Z.F., and Wang, L. (2020). High fat diet-triggered non-alcoholic fatty liver disease: A review of proposed mechanisms. Chem Biol Interact 330, 109199. 10.1016/j.cbi.2020.109199.

33. Caldwell, S.H., Swerdlow, R.H., Khan, E.M., Iezzoni, J.C., Hespenheide, E.E., Parks, J.K., and Parker, W.D., Jr. (1999). Mitochondrial abnormalities in non-alcoholic steatohepatitis. J Hepatol 31, 430–434. 10.1016/s0168-8278(99)80033-6.

34. Bucher, S., Le Guillou, D., Allard, J., Pinon, G., Begriche, K., Tete, A., Sergent, O., Lagadic-Gossmann, D., and Fromenty, B. (2018). Possible Involvement of Mitochondrial Dysfunction and Oxidative Stress in a Cellular Model of NAFLD Progression Induced by Benzo[a]pyrene/Ethanol CoExposure. Oxid Med Cell Longev 2018, 4396403. 10.1155/2018/4396403.

35. Ajith, T.A. (2018). Role of mitochondria and mitochondria-targeted agents in non-alcoholic fatty liver disease. Clin Exp Pharmacol Physiol 45, 413–421. 10.1111/1440-1681.12886.

36. Wei, Y., Rector, R.S., Thyfault, J.P., and Ibdah, J.A. (2008). Nonalcoholic fatty liver disease and mitochondrial dysfunction. World J Gastroenterol 14, 193–199. 10.3748/wjg.14.193.

37. Newman, L.E., and Shadel, G.S. (2023). Mitochondrial DNA Release in Innate Immune Signaling. Annu Rev Biochem 92, 299–332. 10.1146/annurev-biochem-032620-104401.

38. Dornas, W., and Schuppan, D. (2020). Mitochondrial oxidative injury: a key player in nonalcoholic fatty liver disease. Am J Physiol Gastrointest Liver Physiol 319, G400–G411. 10.1152/ajpgi.00121.2020.

39. Al Khatib, I., Deng, J., Lei, Y., Torres-Odio, S., Rojas, G.R., Newman, L.E., Chung, B.K., Symes, A., Zhang, H., Huang, S.N., et al. (2023). Activation of the cGAS-STING innate immune response in cells with deficient mitochondrial topoisomerase TOP1MT. Hum Mol Genet 32, 2422–2440. 10.1093/hmg/ddad062.

40. Pessayre, D. (2007). Role of mitochondria in non-alcoholic fatty liver disease. J Gastroenterol Hepatol 22 Suppl 1, S20–27. 10.1111/j.1440-1746.2006.04640.x.

41. Pessayre, D., Mansouri, A., and Fromenty, B. (2002). Nonalcoholic steatosis and steatohepatitis. V. Mitochondrial dysfunction in steatohepatitis. Am J Physiol Gastrointest Liver Physiol 282, G193–199. 10.1152/ajpgi.00426.2001.

42. Buzzetti, E., Pinzani, M., and Tsochatzis, E.A. (2016). The multiple-hit pathogenesis of non-alcoholic fatty liver disease (NAFLD). Metabolism 65, 1038–1048. 10.1016/j.metabol.2015.12.012.

43. Xu, R., Tao, A., Zhang, S., Deng, Y., and Chen, G. (2015). Association between patatin-like phospholipase domain containing 3 gene (PNPLA3) polymorphisms and nonalcoholic fatty liver disease: a HuGE review and meta-analysis. Sci Rep 5, 9284. 10.1038/srep09284.

44. Kozlitina, J., Smagris, E., Stender, S., Nordestgaard, B.G., Zhou, H.H., Tybjaerg-Hansen, A., Vogt, T.F., Hobbs, H.H., and Cohen, J.C. (2014). Exome-wide association study identifies a TM6SF2 variant that confers susceptibility to nonalcoholic fatty liver disease. Nat Genet 46, 352–356. 10.1038/ng.2901.

45. Liu, Y.L., Reeves, H.L., Burt, A.D., Tiniakos, D., McPherson, S., Leathart, J.B., Allison, M.E., Alexander, G.J., Piguet, A.C., Anty, R., et al. (2014). TM6SF2 rs58542926 influences hepatic fibrosis progression in patients with non-alcoholic fatty liver disease. Nat Commun 5, 4309. 10.1038/ncomms5309.

46. Zhang, H., Seol, Y., Agama, K., Neuman, K.C., and Pommier, Y. (2017). Distribution bias and biochemical characterization of TOP1MT single nucleotide variants. Scientific reports 7, 8614. 10.1038/s41598-017-09258-2.

47. Arendt, B.M., Comelli, E.M., Ma, D.W., Lou, W., Teterina, A., Kim, T., Fung, S.K., Wong, D.K., McGilvray, I., Fischer, S.E., and Allard, J.P. (2015). Altered hepatic gene expression in nonalcoholic fatty liver disease is associated with lower hepatic n-3 and n-6 polyunsaturated fatty acids. Hepatology 61, 1565–1578. 10.1002/hep.27695.

48. Al Khatib, I., Deng, J., Symes, A., Kerr, M., Zhang, H., Huang, S.N., Pommier, Y., Khan, A., and Shutt, T.E. (2022). Functional characterization of two variants of mitochondrial topoisomerase TOP1MT that impact regulation of the mitochondrial genome. J Biol Chem 298, 102420. 10.1016/j.jbc.2022.102420.

49. Kleiner, D.E., Brunt, E.M., Van Natta, M., Behling, C., Contos, M.J., Cummings, O.W., Ferrell, L.D., Liu, Y.C., Torbenson, M.S., Unalp-Arida, A., et al. (2005). Design and validation of a histological scoring system for nonalcoholic fatty liver disease. Hepatology 41, 1313–1321. 10.1002/hep.20701.

50. Kietzmann, T. (2017). Metabolic zonation of the liver: The oxygen gradient revisited. Redox Biology 11, 622–630. 10.1016/j.redox.2017.01.012.

51. Kao, C.Y., Factor, V.M., and Thorgeirsson, S.S. (1996). Reduced growth capacity of hepatocytes from c-myc and c-myc/TGF-alpha transgenic mice in primary culture. Biochem Biophys Res Commun 222, 64–70. 10.1006/bbrc.1996.0698.

52. Gu, Z. (2022). Complex heatmap visualization. Imeta 1, e43. 10.1002/imt2.43.

