## Supplementary Data for "Mitochondrial topoisomerase I (Top1MT) prevents the onset of metabolic dysfunction-associated steatohepatitis (MASH) in mice"

**Supplemental Figures**

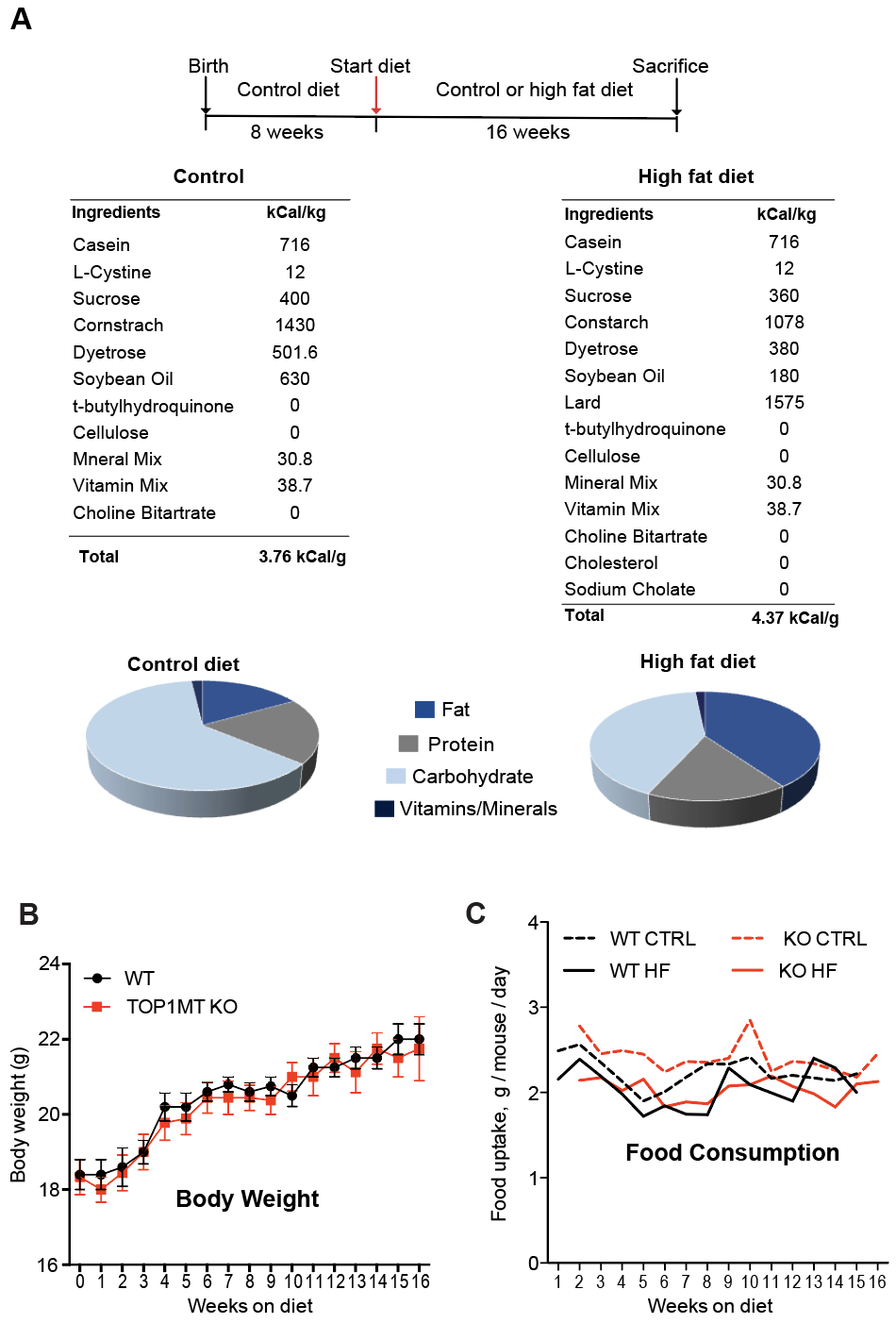

**Figure S1. Dietary model for the analyses of mice and their body wight and food uptake capacity**

(A) Top: Experimental scheme of the diet given to mice and downstream analysis after their sacrifice. Bottom: Components and calories for control and high fat (HF) diet.
(B) Graph representing the gain of body weight in WT and Top1MT KO mice after HF diet consumption for 16 weeks. Body weight was measured in every week.
(C) Graph showing the amount of food uptake in WT and Top1MT KO mice after HF diet consumption for 16 weeks.

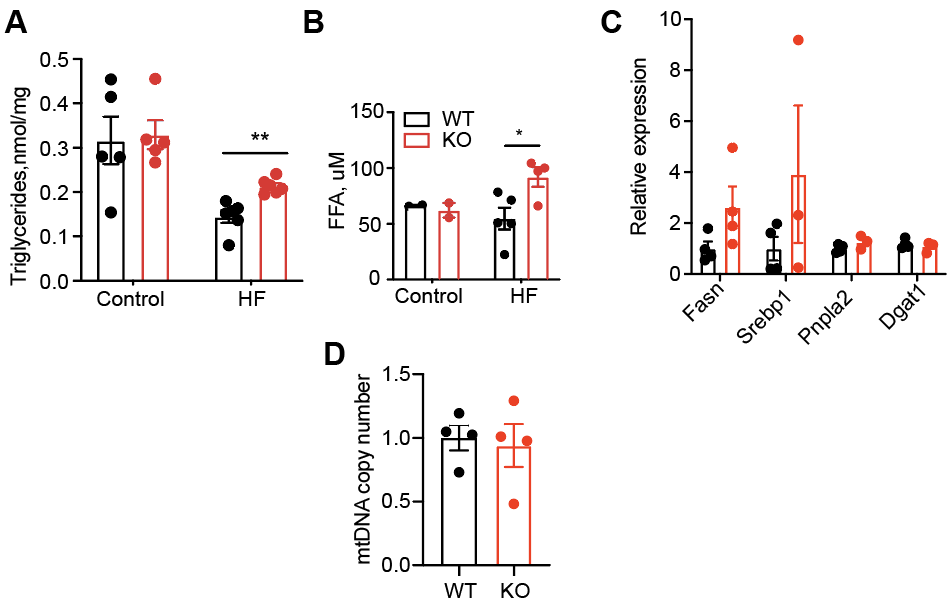

**Figure S2. Altered fat metabolism in Top1MT KO mice after HF diet for 16 weeks**

(A) Measurement of plasma triglycerides levels after 16 weeks on HF diet in WT and Top1MT KO mice. Control, (WT: n=5; KO: n=5) or HF diet, (WT: n=6; KO n=6).
(B) Measurement of plasma free fatty acid (FFA) levels after 16 weeks on HF diet in WT and Top1MT KO mice. Control, (WT: n=2; KO: n=2) or HF diet, (WT: n=5; KO n=4).
(C) Expression level of genes related to fat metabolism were quantified by RT-qPCR in liver tissue after 16 weeks on HF diet. n=3 for both WT and Top1MT KO.
(D) mtDNA copy number was determined by RT-qPCR in adipose tissue from WT and Top1MT KO mice after 16 weeks on HF diet. n=4 for WT and KO both.

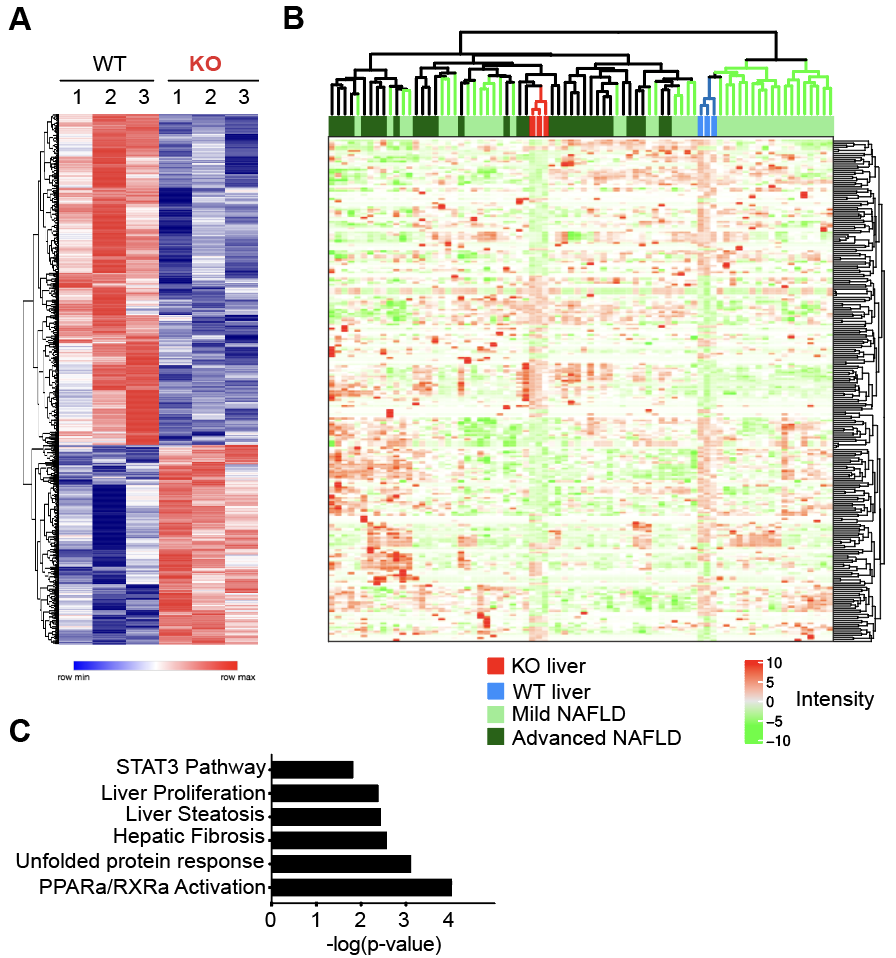

**Figure S3. Top1MT KO gene expression signature predicts disease stage of NAFLD patients**

(A) Heatmap of significant differentially expressed genes in liver tissues of WT and Top1mt KO mice after 16 weeks on HF diet. n=3 mice for both WT and KO.
(B) Integrative cluster analysis of murine MASH gene signature applied to NAFLD patients using orthologous genes. n=3 mice, KO liver; n=3 mice, WT liver; n=40 patients, Mild NAFLD and n=32 patients, Advanced NAFLD.
(C) KEGG pathway analysis of significantly altered genes.
